# Rational identification of potent and broad sarbecovirus-neutralizing antibody cocktails from SARS convalescents

**DOI:** 10.1101/2022.08.03.499114

**Authors:** Yunlong Cao, Fanchong Jian, Zhiying Zhang, Ayijiang Yisimayi, Xiaohua Hao, Linlin Bao, Fei Yuan, Yuanling Yu, Shuo Du, Jing Wang, Tianhe Xiao, Weiliang Song, Ying Zhang, Pulan Liu, Ran An, Peng Wang, Yao Wang, Sijie Yang, Xiao Niu, Yuhang Zhang, Qingqing Gu, Fei Shao, Yaling Hu, Weidong Yin, Aihua Zheng, Youchun Wang, Chuan Qin, Ronghua Jin, Junyu Xiao, Xiaoliang Sunney Xie

## Abstract

SARS-CoV-2 Omicron sublineages have escaped most RBD-targeting therapeutic neutralizing antibodies (NAbs), which proves the previous NAb drug screening strategies deficient against the fast-evolving SARS-CoV-2. Better broad NAb drug candidate selection methods are needed. Here, we describe a rational approach for identifying RBD-targeting broad SARS-CoV-2 NAb cocktails. Based on high-throughput epitope determination, we propose that broad NAb drugs should target non-immunodominant RBD epitopes to avoid herd immunity-directed escape mutations. Also, their interacting antigen residues should focus on sarbecovirus conserved sites and associate with critical viral functions, making the antibody-escaping mutations less likely to appear. Following the criteria, a featured non-competing antibody cocktail, SA55+SA58, is identified from a large collection of broad sarbecovirus NAbs isolated from SARS convalescents. SA55+SA58 potently neutralizes ACE2-utilizing sarbecoviruses, including circulating Omicron variants, and could serve as broad SARS-CoV-2 prophylactics to offer long-term protection. Our screening strategy can also be applied to identify broad-spectrum NAb drugs against other fast-evolving viruses, such as influenza viruses.

## Main

Over two years after its emergence, the COVID-19 pandemic caused by SARS-CoV-2 is still spreading. Neutralizing antibodies (NAbs) play a critical role in the prevention and treatment of COVID-19^1-4^. However, the constant emergence of new variants has caused large-scale evasion of NAbs, posing severe challenges to SARS-CoV-2 NAb drugs^5-9^. Given its prophylactic and therapeutic efficacy, the clinical development of NAb drugs is still in high demand^10,11^, especially those broad SARS-CoV-2 NAbs that are difficult for future variants to escape.

All currently approved SARS-CoV-2 NAb drugs target the receptor-binding domain (RBD) of the SARS-CoV-2 spike glycoprotein, such as LY-CoV016+LY-CoV555 (bamlanivilab+etesevimab)^12,13^, REGN10933+ REGN10987 (casirivimab+imdevimab)^10,14^, S309 (sotrovimab)^15^, AZD1061+AZD8895 (cilgavimab+tixagevimab, AZD7442, Evusheld)^16^ and LY-CoV1404 (bebtelovimab)^17^. Unfortunately, the majority of them have already been escaped by the Omicron variants^6,8,18,19^. Most of the NAbs were selected for clinical development mainly based on two criteria: I) The antibody candidates were potent against the circulating variants at the time^10,12-17,20^. II) The antibody candidates should better form a non-competing antibody cocktail^10,16,21^. Given that high potency could lead to lower dosage and non-overlapping cocktails would reduce the chance of being completely escaped, the strategy is reasonable; however, it is insufficient. Practically, the virus was able to evolve multiple mutations on the RBD that could escape both antibodies in a cocktail simultaneously^22,23^, and high potency would not guarantee good neutralization breadth against SARS-CoV-2. Better broad NAb drug selection strategies, beyond simply picking the most potent ones to form cocktails, are needed to counter the fast-evolving virus.

One potential broad SARS-CoV-2 NAb drug selection strategy is to choose NAbs that exhibit broad sarbecovirus neutralizing activity^3,4^. These broad sarbecovirus SARS-CoV-2 NAbs (bsNAbs) have epitopes located on sarbecovirus conserved regions, and people believe that they would not be easily escaped by SARS-CoV-2 variants. Various bsNAbs targeting spike protein’s RBD or S2 region have been discovered^24-27^. S2-targeting bsNAbs indeed displayed exceptional SARS-CoV-2 neutralization breadth, but their potency is rather low for NAb drug development. On the other hand, anti-RBD bsNAbs, such as S309^15^, ADG-2^28^, DH1047^29^, S2X259^4^, and S2K146^30^, could exhibit high neutralization potency and are good candidates for NAb drugs; however, the majority of them are escaped by Omicron BA.2 or BA.4/BA.5^6,18^. This suggests that NAbs with broad sarbecovirus neutralizing capability are actually not directly equivalent to broad SARS-CoV-2 NAbs, and a better broad NAb drug selection strategy is still needed.

In this paper, we describe a rational approach for identifying RBD-targeting broad SARS-CoV-2 NAb cocktails with high potency that are strongly resistant to both current and potential future mutants. The resulting featured antibody cocktail, SA55+SA58, exhibits broad sarbecovirus neutralizing activity and is highly potent against current Omicron variants, including BA.1, BA.2, BA.2.12.1 and BA.4/BA.5, making it a valuable bsNAb drug candidate.

## Results

### A rational strategy for identifying potent bsNAb cocktails

We believe that rational identification of broad SARS-CoV-2 NAb drug candidates relies on the accurate estimation of antibodies’ neutralization breadth; however, neutralization breadth is not only governed by the biochemical properties of the NAbs but also depends on the virus mutation pattern and evolution direction. Based on the study of SARS-CoV-2 variants, the evolution of the virus RBD seems to comply with the following patterns: SARS-CoV-2 is more likely to evolve RBD mutations on “hot” epitopes that are frequently targeted by NAbs elicited by SARS-CoV-2 convalescents and vaccinees to achieve efficient humoral immunity evasion^6,23,31^; also, the virus is less likely to evolve mutations that could disrupt its conserved functions, such as ACE2-binding or RBD-folding^32^. On the basis of these observations, we propose two additional criteria for identifying RBD-targeting broad SARS-CoV-2 NAb drugs: III) The candidate NAbs should target a rare RBD epitope, such that NAbs with similar epitope are not abundantly elicited by SARS-CoV-2 convalescents and vaccinees. This way, the RBD mutations that escape the candidate NAbs are not aligned with those that heavily evade herd humoral immunity and would have a lower chance of appearing. IV) The candidate NAbs should target sarbecovirus conserved epitopes on the RBD that are also associated with critical viral functions, such as ACE2-binding or glycosylation. These NAbs would naturally exhibit broad sarbecovirus binding capability, and their escaping mutations are less likely to prevail^30^.

However, several difficulties exist for identifying NAb candidates that meet these four criteria. First, a substantially large antibody library is almost mandatory to contain a collection of rare anti-RBD NAbs that target sarbecovirus conserved regions. To reduce the size of the antibody library needed, we choose SARS-CoV-2 vaccinated SARS convalescents as the antibody source since their memory B cells are more likely to encode NAbs that target sarbecovirus conserved epitopes^33^. Second, a high-throughput method is needed to determine the binding epitopes and antibody-escaping mutations of each NAb in a large library. Previously, we presented a high-throughput unsupervised epitope mapping technology based on deep mutational scanning (DMS), which can be employed for epitope distribution and escaping mutation analyses at much higher efficiency than traditional epitope binning techniques^23,34^. By combining high-throughput DMS with droplet-based single-cell V(D)J sequencing (scVDJ-seq) or single-cell reverse transcription PCR (RT-PCR) of RBD-specific memory B cells^20,21,35,36^, an efficient pipeline can be built to isolate a large panel of NAbs and, at the same time, identify their binding epitopes, solving the above difficulties^6^ (Fig. 1a).

**Fig. 1.**
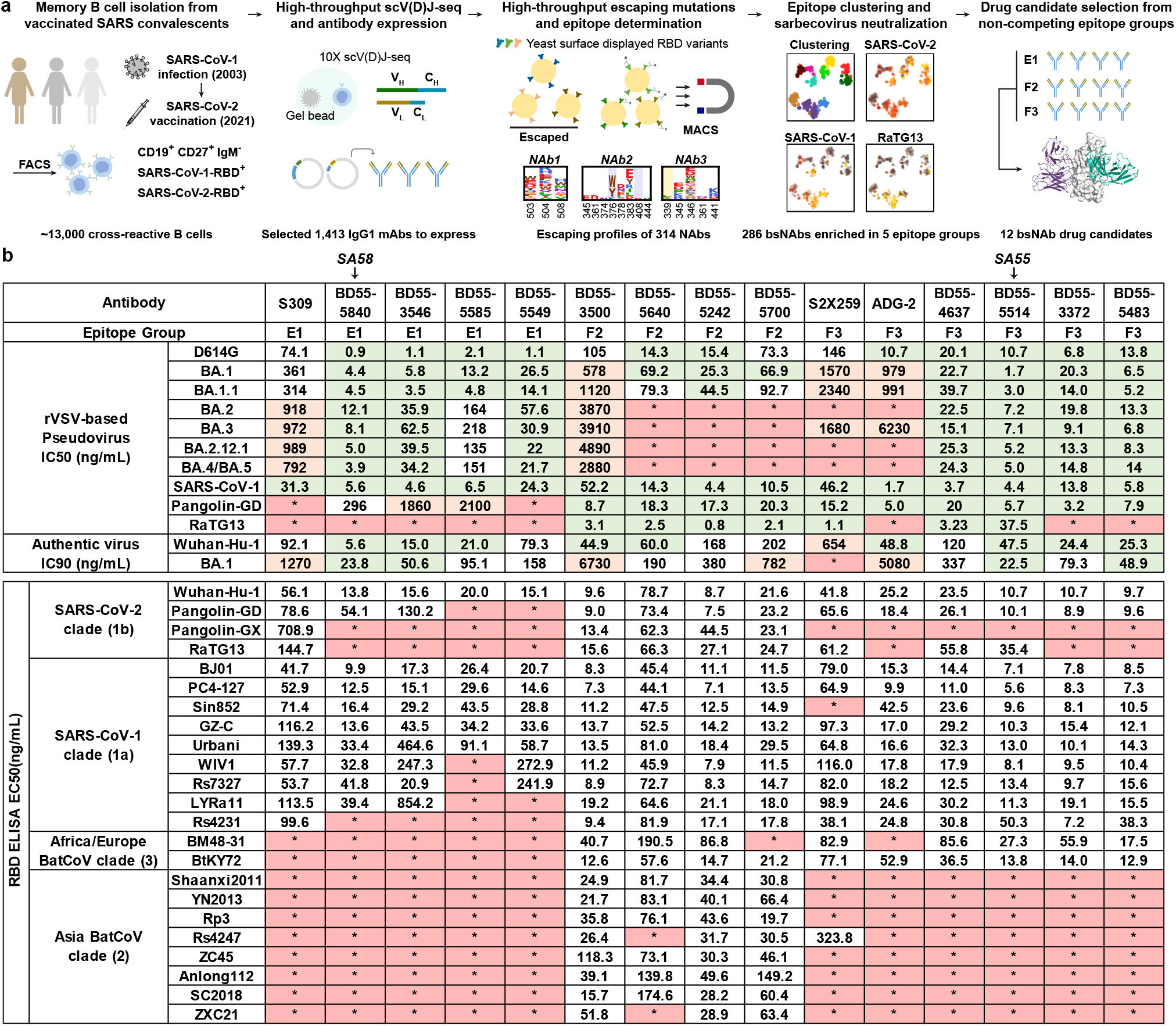
Identification of broad sarbecovirus neutralizing antibodies. a, Schematic for the identification and characterization of bsNAbs, and selection of bsNAb drug candidates from SARS-CoV-2-vaccinated SARS convalescents. b, Neutralizing IC50 against SARS-CoV-2 variants VSV-based pseudoviruses and authentic viruses, and ELISA EC50 against sarbecovirus RBD by identified bsNAbs and other published antibodies in group E1, E3 and F3.

As proposed, we successfully obtained a large bsNAb collection from SARS-CoV-2 vaccinated SARS convalescents, with each antibody’s epitope and escaping mutational profile determined (Fig. 1a). Specifically, we used fluorescence-activated cell sorting (FACS) and collected ∼13,000 CD19^+^CD27^+^IgM^-^ memory B cells that can cross-bind to both SARS-CoV-2 RBD and SARS-CoV-1 RBD from 28 individuals who recovered from SARS in 2003 and received 2 doses of SARS-CoV-2^WT^ inactivated vaccine (CoronaVac) and 1 booster dose of RBD-based protein subunit vaccine (ZF2001) in 2021^37,38^ (Extended Data Fig. 1, Supplementary Table 1). A total of 2838 heavy-light chain paired antibody sequences were recovered from those cross-reactive memory B cells by high-throughput scVDJ-seq^20^. Among them, 1413 antibody sequences contain IgG1 heavy chain constant region, and were selected to express *in vitro* as monoclonal antibodies (mAbs) (Supplementary Table 2). The neutralizing activity of the mAbs against SARS-CoV-2 was further screened using VSV-based pseudovirus (D614G). We then selected NAbs (D614G IC50 < 10μg/ml) for further epitope mapping and successfully obtained the RBD escaping mutational profiles of 314 NAbs, among which 286 displayed sarbecovirus neutralizing activity (SARS-CoV-1 IC50 < 10μg/ml)^6,23^ (Fig. 1a, Supplementary Table 2).

These bsNAbs mainly appeared in five epitope groups (E1, E3, F1, F2 and F3), determined by unsupervised clustering based on DMS results as previously presented^6,23^ (Extended Data Fig.2). In general, Group E1 (S309 epitope^15^) antibodies bind to the front of RBD and exhibit the binding capability to all clade 1a/1b sarbecoviruses (Fig. 1b). F3 (ADG-2 epitope^28^) NAbs bind to the back of RBD and could bind to all ACE2-utilizing sarbecovirus clades, including BtKY72 and BM48-31 in clade 3 (Fig. 1b). Group E3 (S2H97^27^ epitope), F1 (S304^15^ epitope) and F2 (BD55-1239^6^ epitope) antibodies exhibited broad specificity to sarbecoviruses in all clades (Fig. 1b); however, antibodies from cryptic epitope E3 and F1 exhibited low neutralizing activities, and are not suitable for NAb drugs. In comparison, E1 and F3 NAbs displayed high neutralizing activity while F2 demonstrated moderate potency, but better breadth (Fig. 1b). A total of 49 E1, 109 F2, and 57 F3 bsNAbs were identified (Supplementary Table 2).

According to criterion I, we selected the top four most potent NAbs from each of the three epitope groups, E1, F2 and F3, as bsNAb drug candidates based on the pseudovirus neutralizing activity against SARS-CoV-1, SARS-CoV-2 D614G, and Omicron BA.1, which is the dominant variant at the time. The majority of bsNAb candidates showed high somatic hypermutation rates, which resulted from the long-term antibody affinity maturation since SARS infection in 2003 (Extended Data Fig. 3). Among the four candidates in Group E1, BD55-5840 showed the highest authentic virus neutralization activity against both the SARS-CoV-2 ancestral strain (IC90=5.6ng/mL) and Omicron BA.1 (IC90=23.8ng/mL), much stronger than S309, the archetypical E1 antibody (Fig. 1b). In Group F3, BD55-5514 exhibited the highest activity against BA.1 authentic virus (IC90=22.5ng/mL), which is much more potent than ADG-2 (Fig. 1b). The overall authentic virus neutralization of NAbs from Group F2 is indeed much lower than that of E1 and F3 (Fig. 1b).

**Fig. 2.**
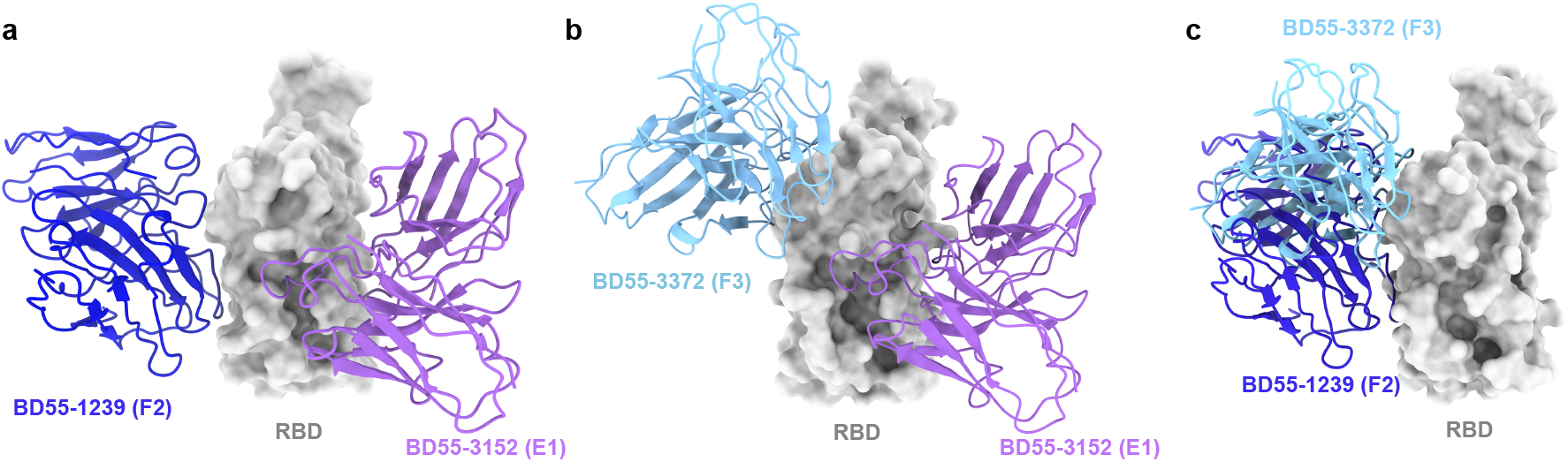
Proposed strategies for non-overlapping bsNAb combination. a-b, Aligned structures of BD55-3152 (Group E1, PDB: 7WR8) together with a, BD55-1239 (Group F2, PDB: 7WRL) or b, BD55-3372 (Group F3, PDB: 7WRO) in complex with SARS-CoV-2 BA.1 RBD. c, Aligned structure shows BD55-1239 (PDB: 7WRL) in F2 and BD55-3372 (PDB: 7WRO) in F3 could not form non-overlapping cocktails.

**Fig. 3.**
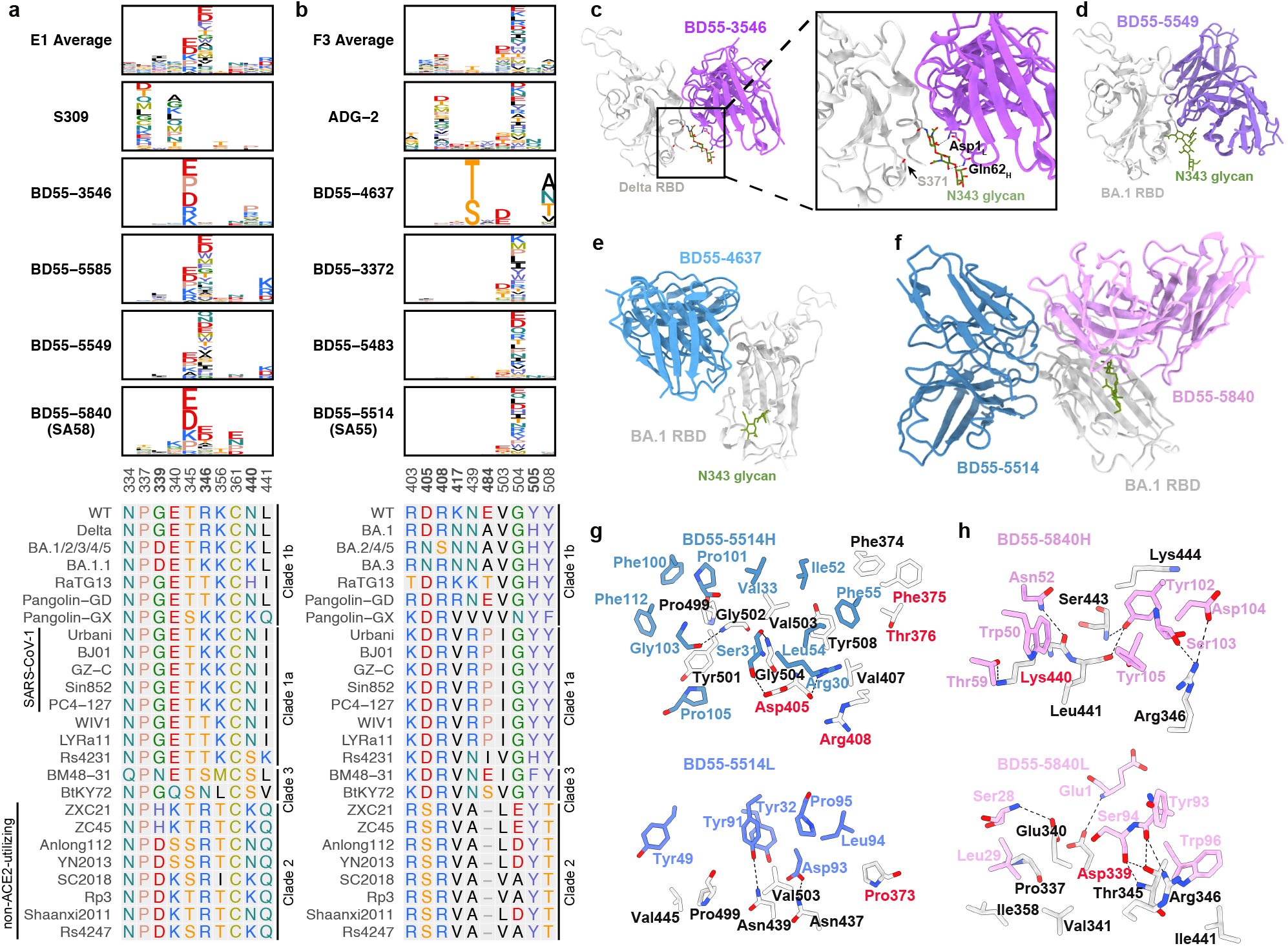
Escape maps and structural analyses of representative E1 and F3 antibodies. a-b, Average escape scores of antibodies in group E1 (a) and F3 (b), and escape maps of selected drug-candidate bsNAbs in group E1 (a) and F3 (b). MSA on the corresponding escaping hotspot residues are shown. c, Structure of BD55-3546 bound to RBD^Delta^. d, Structure of BD55-5549 bound to RBD^BA.1^. e, Structure of BD55-4637 bound to RBDBA.1. f, Structure of BD55-5514 and BD55-5840 bound together to RBD^BA.1^. g, Interactions between BD55-5514 and RBD^BA.1^. h, Interactions between BD55-5840 and RBD^BA.1^.

Next, we examined the possible cocktail pairing strategies among these three epitope groups. We performed antibody structural alignment analyses based on three representative antibody structures, BD55-5840 for E1, BD55-1239 for F2, and BD55-3372 for F3 (Fig. 2a-c)^6^. Results demonstrated that E1 antibodies could generally form non-overlapping cocktails with antibodies in either Group F2 or F3 (Fig. 2a-b), while F2 and F3 antibodies covered overlapping epitopes (Fig. 2c). Thus, E1+F2 and E1+F3 are the two practical strategies to build non-competing bsNAb cocktails. However, we previously showed that F2 antibodies are abundant in SARS-CoV-2^WT^ convalescents/vaccinees, while E1 and F3 are generally rare^6^. Therefore, E1/F3 cocktails are better than E1/F2 cocktails, according to criterion III, such that the rarity of these NAbs would make their escaping RBD mutations less likely to prevail. Also, the epitopes of E1 NAbs are centered around N343 glycan (Fig. 3a), and F3 NAbs’ escaping mutations mainly focus on the binding interface of ACE2 (Fig. 3b), while the epitopes of F2 NAbs are not linked to any known critical viral functions. Indeed, almost all F2 NAbs were escaped by Omicron BA.2 subvariants mainly due to the D405N and R408S locating on their epitopes, despite these two sites being highly conserved among sarbecoviruses (Fig. 1b)^23^. This exemplifies criterion IV, that antibodies targeting sarbecovirus conserved epitopes can also be easily escaped, unless their major escaping mutations directly overlap with sites that associate with critical viral functions and are less likely to mutate.

### Structural analyses of the bsNAb drug candidates

To decide the final bsNAb cocktail composition, we further examined the escaping mutation profiles and structural properties of the E1 and F3 NAb candidates. Most E1 NAbs are susceptible to mutations of T345 and R346 (Fig. 3a). The four E1 bsNAbs displayed three different escaping mutation profiles, suggesting three different RBD-binding modes. BD55-3546 is mainly escaped by T345 and N440 mutations. BD55-5585 and BD55-5549 have similar escaping mutation profiles and are mainly escaped by R346, T345, and L441 mutations. BD55-5840 is less affected by R346 mutations compared to BD55-5585 and BD55-5549. To analyze their detailed molecular interactions with RBD, we determined the crystal structure of BD55-5549/RBD_BA.1_ and the cryo-electron microscopy (cryo-EM) structures of BD55-3546/S6P_Delta_ (Fig. 3c-d, Extended Data Fig. 4a, Supplementary Table 3 and 4). Previously, we also solved the cryo-EM structures of BD55-5840/S6P_BA.1_ and BD55-5840/S6P_BA.2_ ^6^. Similar to S309, the epitopes of these antibodies all encompass the N343 glycan, which plays a critical role in modulating RBD conformation. Thus, critical mutations that can alter the position of the glycan motif could reduce the activities of the E1 antibodies, such as S371F that emerged in BA.2^6^. BD55-3546 is more affected by this mutation than BD55-5840, with 5∼10-fold decreased activities towards the S371F-containing Omicron variants (Fig. 1b). In the BD55-3546/S6P_Delta_ structure, Q62_H_ and D1_L_ pack intimately with the N343 glycan (Extended Data Fig. 5a), rationalizing the sensitivity of BD55-3546 to its displacement. BD55-3546 is also sensitive to mutations of N440, as N440 is nestled in a pocket formed by W50_H_, N52_H_, T55_H_, I57_H_, and T59_H_ and forms extensive van der Waals and hydrogen bond interactions with these residues. By comparison, BD55-5549 is not affected by the changes of N440. Indeed, the N440K site is not involved in interacting with BD55-5549 (Extended Data Fig. 5b). On the other hand, L441 is surrounded by Y110_H_, Y110_H_, and F111_H_, and BD55-5549 could be escaped by the L441 mutations (Fig. 3a). Contrarily, BD55-5840 is not susceptible to the substitutions of 440-441, as the side chain of neither site is closely targeted by BD55-5840; and is only slightly affected by S371F^6^. Mutations of T345 could lead to prominent escapes from BD55-5840 (Fig. 3a), since T345 is enclosed by Y105_H_, L94_L_, and W96_L_; however, T345 substitutions are unlikely to emerge, as it is essential for the proper glycosylation of N343 that is critical for RBD function^32^. Together, considering the prevalence of R346 (BA.1.1) and S371 (BA.2) mutations, we selected BD55-5840 as the final E1 bsNAb for clinical development due to its higher potency and lesser sensitivity to glycan displacement and R346 substitutions.

**Fig. 4.**
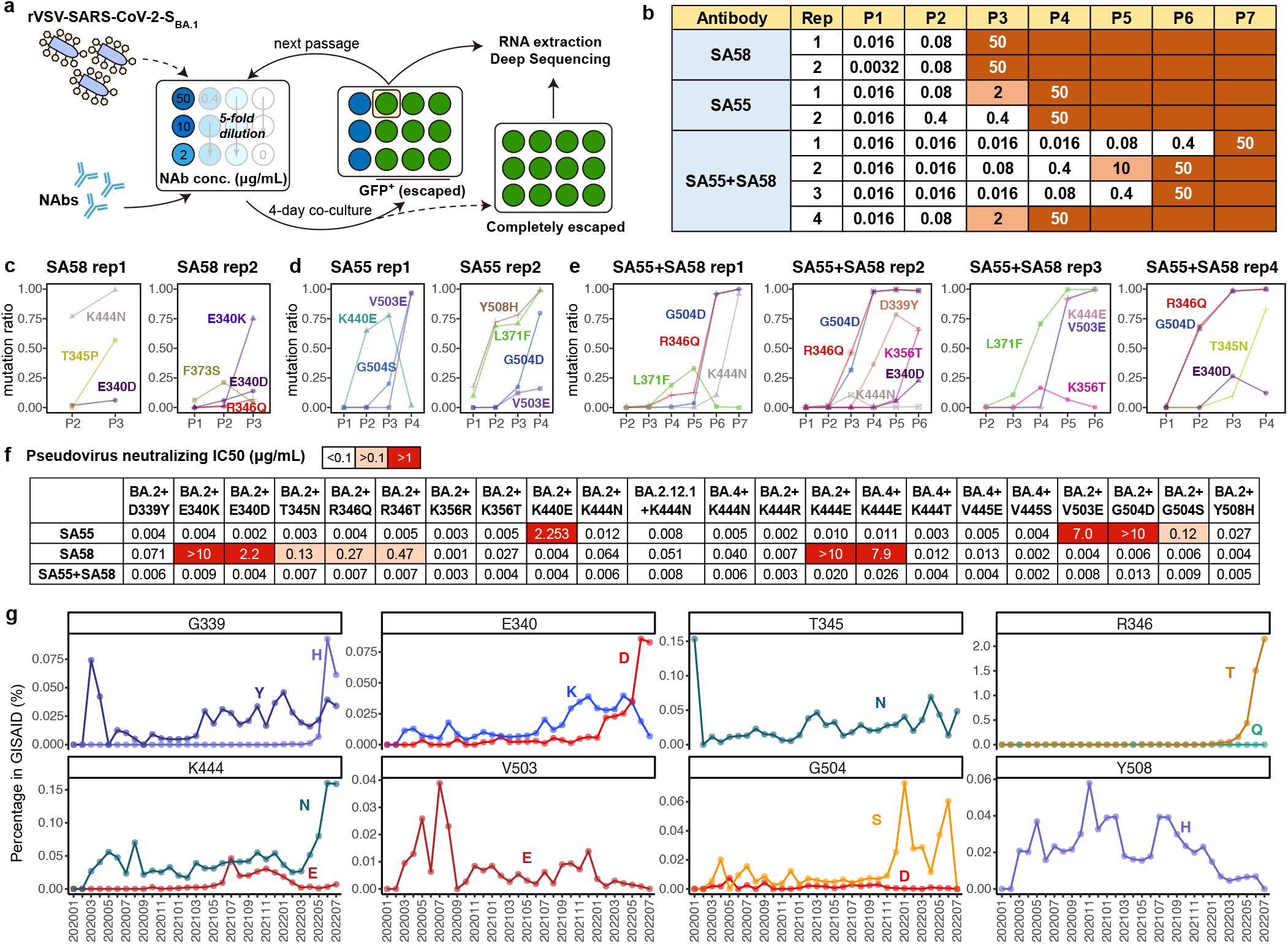
SA55/SA58 cocktail robustly neutralizes SARS-CoV-2 escaping mutants. a, Schematic of in vitro escape mutation screening using SARS-CoV-2^BA.1^-spike-pseudotyped replication-competent recombinant VSVs (rVSV). The virus was passed in the presence of antibody dilutions for 4 days on Vero cells. Cells were screened for virus replication by monitoring GFP+. b, Antibody concentration of dilutions that are escaped, passed, and sequenced of each passage and each antibody. SA55 and SA58 alone were tested in two biological replicates, respectively. SA55+SA58 cocktail was tested in four biological replicates. c-e, The ratio of enriched escape mutations of each replicate and each passage by deep sequencing. c, using SA58 only; d, using SA55 only; e, SA55+SA58 cocktail. f, Neutralizing activities by SA55, SA58 and SA55+SA58, against selected SARS-CoV-2 variants harboring potential escaping mutations of SA55 or SA58. g, Ratio of selected SA55/SA58-escaping mutations in detected SARS-CoV-2 sequences from GISAID.

**Fig. 5.**
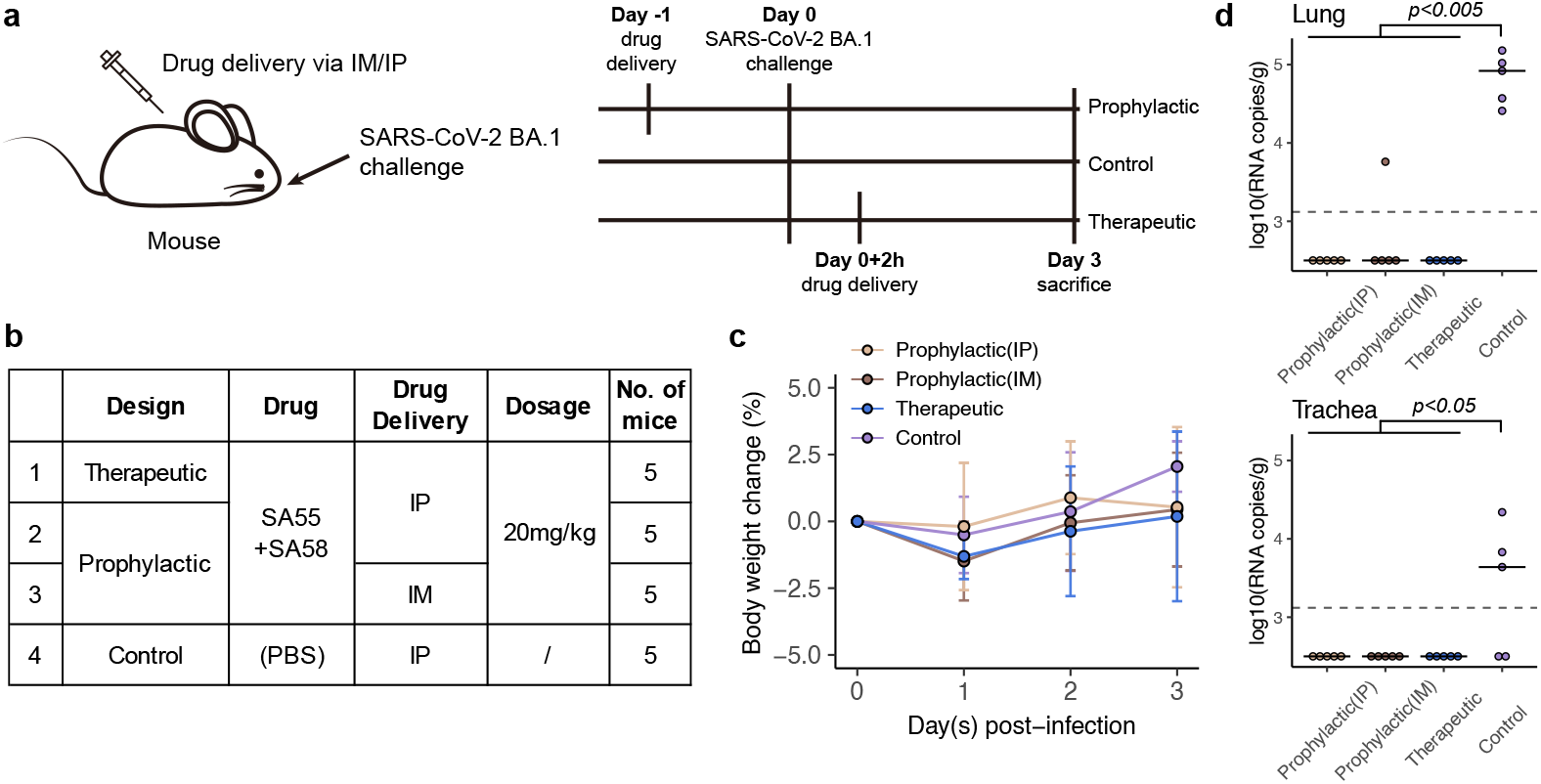
SA55/SA58 demonstrate potent protection efficacy in mice. a-b, Design for the SARS-CoV-2 BA.1 challenge experiment in mice. In prophylactic groups, 20mg/kg SA55+SA58 were administered intraperitoneally or intramuscularly and challenged by SARS-CoV-2 BA.1 intranasally one day later. In therapeutic groups, antibodies were given intraperitoneally 2 hours after virus challenge. Each group consists of 5 mice. c, Percentage of changes in body weight compared to the weight of each mouse before the experiment. Error bars indicate mean±s.d. d, Viral load in lung (up) and trachea (bottom) of mice in each group. Each point corresponds to samples from a mouse. P-values were calculated using one-tailed Wilcoxon rank-sum test. The alternative hypothesis is that the values in prophylactic or therapeutic groups are less than values in control groups.

F3 NAbs are mainly escaped by mutations of V503 and G504, as well as the D405 mutation found in BA.2 (Fig. 3b). BD55-4637 has a rather unique escaping mutation profile, while BD55-3372, BD55-5483 and BD55-5514 share highly similar profiles, suggesting similar interactions with RBD (Fig. 3b). BD55-4637 can be escaped by the N439T/S mutations, as these mutations would lead to the glycosylation of N437, which packs against W55_H_, revealed by the cryo-EM structure of BD55-4637/S6P_BA.1_ (Fig. 3e, Extended Data Fig. 4b, Extended Data Fig. 5c). BD55-4637 is also sensitive to mutations of Y508, due to the interaction between Y508 and D102_H_; however, Y508 is essential to ACE2 binding and could hardly be evolved^32^ (Extended Data Fig. 5c). In contrast, BD55-3372, BD55-5483, and BD55-5514 are not susceptible to the changes of these two sites, but are sensitive to the changes on V503 and G504 (Fig. 3b). Nevertheless, V503 and G504 are also critical for ACE2 binding as showed by deep mutational scanning (DMS) and are conserved among sarbecoviruses^32^, making them difficult to prevail. Based on these characteristics, all four F3 bsNAbs are good drug candidates since their escaping mutations could not easily appear, and BD55-5514 is chosen as the final F3 bsNAb for clinical development due to its highest neutralization potency (Fig. 2b).

BD55-5514 indeed can bind to the RBD together with BD55-5840 in a largely non-interfering manner, as demonstrated by the BD55-5840/BD55-5514/S6P_BA.1_ ternary complex structure (Fig. 3f, Extended Data Fig. 4c). Interestingly, a close examination of the BD55-5514/S6P_BA.1_ structure reveals that the S373P and S375F mutations could promote the interaction with BD55-5514, consistence with the pseudovirus results (Fig. 2b). F374 is flipped out because of these two mutations and robustly packs with F55_CDRH2_ in BD55-5514 (Fig. 3g). P373_BA.1_ also forms a hydrophobic interaction with L94_CDRL3_ (Fig. 3g). Although T376, D405, and R408 are involved in the interaction with BD55-5514, they are all located at the periphery of BD55-5514’s epitope, and the activities of BD55-5514 towards BA.2/BA.2.12.1/BA.2.13/BA.4/BA.5 are only slightly decreased when compared to BA.1 (Fig. 2b, Fig. 3g). Most importantly, BD55-5840 and BD55-5514 could synergize in a complementary way. BD55-5840 does not directly impede ACE2 but can engage both the up and down RBDs, whereas BD55-5514 only binds to the up RBDs but promptly blocks ACE2 (Extended Data Fig. 6). Together, BD55-5840 (SA58) + BD55-5514 (SA55) is rationally selected as the final non-competing bsNAb cocktail, which exhibits exceptional SARS-CoV-2 neutralization breadth and potency.

### SA55/SA58 display broad resilience against RBD single substitutions

To further explore possible escape mutants of SARS-CoV-2 against SA55/ SA58, we screened for escape mutations by infesting Vero cells with replication-competent recombinant VSV (rVSV)-SARS-CoV-2-S_BA.1_ at a gradient of neutralizing antibody concentration to simulate the evolutionary stress caused by NAbs^39,40^. After four days of co-culture with different concentrations of antibodies and rVSV-SARS-CoV-2-S^BA.1^, GFP^+^ wells reflect significant viral replication and antibody evasion (Fig. 4a). The supernatant and cell layer of the well that has the highest antibody concentration among the GFP^+^ wells were collected. Total RNA was extracted and subjected to reverse transcription, amplification, and deep sequencing. Next-generation sequencing reads were aligned to codon-optimized Omicron BA.1 spike nucleotide sequence to determine enriched substitutions (Fig. 4a).

In both replicates, SA58 was escaped in Passage 3 (P3), but the evasion could be attributed to different mutations (Fig. 4b). In rep1, SA58 showed a decreased activity in P2 due to K444N, and completely escaped by the combination of K444N and T345P in P3 (Fig. 4c). Compared to rep1, the evasion of SA58 in rep2 was dominated by mutations on E340 instead of K444. Specifically, a slight enrichment of E340D in P2 resulted in a mild drop in potency, and the sharp enrichment of E340K in P3 completely escaped SA58 (Fig. 4c). Slightly better than using SA58 alone, SA55 alone was not escaped until P4 in both replicates. In SA55 rep1, K440E was enriched in P2 and P3, but disappeared in P4 when it was replaced by G504S and V503E (Fig. 4d). In SA55 rep2, the enrichment of Y508H in P2/P3 resulted in a drop of activity, and the final evasion was caused by G504D/V503E in P4, similar to rep1. Interestingly, L371F was identified since P2 in SA55 rep2, which is far from its binding interface and not an escaping mutation against SA55, given that SA55 could potently neutralize BA.2, which harbors 371F. This may be due to the potential advantage of 371F in cell infection compared to 371L.

In comparison with using SA55 or SA58 alone, inhibition by the SA55+SA58 cocktail could survive more passages, ranging from P4 to P7 in four replicates (Fig.4b). In rep1, the neutralization by SA55+SA58 cocktail was not affected until the co-occurrence of G504D (which escaped SA55) and R346Q (which affected SA58) in P6, and the remaining activity of SA58 was finally eliminated by K444N in P7. Co-occurrence of G504D and R346Q was also observed at rep2 in P3-P4 and rep4 in P2-P3, while the remaining activity of SA58 was eliminated by D339Y/E340D in rep2 and T345N/E340D in rep4, respectively (Fig. 4e). In rep3, L371F was observed again, which slightly affected the binding affinity of SA58, as reported previously^6^. The final evasion was caused by the combination of SA58-escaping K444E and SA55-escaping V503E. These results demonstrate the enhanced ability of SA55+SA58 cocktail to resist escaping mutations compared to their individual usage. Similar to most existing antibody cocktails, at least two escaping substitutions are required to escape SA55+SA58.

To further validate the identified escaping mutations and the broad activity of SA55+SA58 cocktail, we constructed multiple VSV-based pseudoviruses harboring SARS-CoV-2 BA.2/BA.2.12.1/BA.5 spike with additional substitutions on RBD, including 339Y, 340K/D, 345N, 346Q/T, 356R/T, 440E, 444N/R/E/T, 445E/S, 503E, 504D/S, or 508H, and determined the neutralizing activity of SA55, SA58 and SA55+SA58 cocktail against them. The results correspond well with the above DMS, structural analyses and rVSV-based mutation screening. Specifically, the efficacy of SA55 was slightly affected by Y508H, moderately affected by G504S, strongly affected by K440E, and escaped by V503E and G504D. And SA58 was slightly affected by K356T, D339Y and K444N, moderately affected by T345N, R346Q and R346T, strongly affected by E340D, and escaped by E340K and K444E (Fig. 4f and Extended Data Fig. 7). However, most of the identified escaping mutations would lead to changes in surface electricity, locate on sites that are conserved across ACE2-utilizing sarbecoviruses (Fig. 3a-b), and have not been circulating before (Fig. 4g). A notable exception is R346T which is not conserved and has been emerging recently. Fortunately, R346T only causes a moderate drop in SA58 activity and does not affect SA55. SA55+SA58 could potently neutralize all constructed SARS-CoV-2 Omicron mutants, demonstrating its noteworthy breadth and resistance against escaping mutations.

### SA55/SA58 demonstrate high viral clearance efficacy *in vivo*

To evaluate the therapeutic and prophylactic efficacy of SA55+SA58 cocktail against SARS-CoV-2 *in vivo*, human ACE2 (hACE2)-transgenic mice were challenged by infectious SARS-CoV-2 BA.1 isolate before or after treatment by SA55+SA58. In prophylactic and therapeutic groups, 20 mg/kg of 1:1 SA55 and SA58 was given to the hACE2-transgenic mice via intraperitoneal (IP) injection 24 h pre-infection or 2 h post-infection, respectively (Fig. 5a-b). To test the robustness of different antibody delivery approaches, an additional prophylactic group of mice received SA55+SA58 via intramuscular (IM) injection. A control group of mice that received phosphate-buffered saline (PBS) instead of SA55+SA58 was also included. Each group consists of five mice (Fig. 5b). The body weight of each mouse in each group was monitored and recorded daily (Fig. 5c). All of the mice were sacrificed at day 3, and the lungs and trachea were collected for viral load analyses. In general, mice in the control group exhibited significantly higher lung and trachea viral RNA copies compared to those in all prophylactic and therapeutic groups. Specifically, all of the five mice in the control group exhibited a viral load of >10^4^ RNA copies per gram in lung. However, in trachea, virus was only detected in three of the five mice in the control group (Fig. 5d). In contrast, except that one mouse in the prophylactic (IM) group, virus RNA was not detected in lungs and trachea of all mice in all prophylactic or therapeutic groups. Notably, BA.1-infected mice only displayed mild diseases that do not lead to obvious changes in body weight (Fig. 5c). Together, SA55+SA58 displayed strong viral clearance and blocking efficacy *in vivo*.

## Discussion

In this study, we proposed a rational strategy to discover broad NAb drugs that are strongly resistant to SARS-CoV-2 RBD mutations. We hypothesized that a good broad NAb candidate should avoid targeting the public immunodominant epitopes, such that the virus will less likely evolve mutations on the susceptible residues of these NAb drug candidates; also, the drug candidates should target conserved residues that are associated with critical functions, such as receptor binding or antigen folding, which will enable stronger resistance to viral mutations. By integrating high-throughput DMS with scVDJ-seq, we successfully obtained a large collection of bsNAbs from SARS-CoV-2-vaccinated SARS convalescents, and rationally selected a pair of featured bsNAbs, SA55/SA58, to form a non-competing cocktail that demonstrated exceptional neutralizing breadth and potency against SARS-CoV-2 VOCs and related sarbecoviruses. Also, SA55+SA58 exhibited strong resistance to RBD mutations and are potentially difficult to be escaped by future SARS-CoV-2 variants. We believe this rational strategy can also be applied to discover broadly reactive antibody drugs against other rapidly evolving viruses.

Due to antibodies’ prolonged half-life, NAb drugs could serve as long-term prophylactics, where small molecule drugs, such as Paxlovid, could not^41,42^. When engineered on the Fc region, such as YTE modification, the half-life of NAbs can be further extended to near 90 days, suggesting that NAb drugs can stay effective for around half a year^41^. Therefore, Fc-engineered SA55+SA58 are designed to serve as long-term broad SARS-CoV-2 prophylactic NAb drugs, similar to Evusheld, which has been approved to be used as pre-exposure prophylaxis for the prevention of SARS-CoV-2 symptomatic infections. Compared to Evusheld, SA55+SA58 is more difficult for SARS-CoV-2 to escape, making it an advantageous drug candidate to afford long-lasting protection against current and future sarbecovirus mutants. SA55+SA58 would be especially valuable for elderly and immunocompromised individuals who cannot produce enough effective antibodies or are not suitable for vaccination. SA55+SA58 is scheduled to enter phase 1 clinical trial in December 2022.

## Data Availability

Processed rVSV-based mutant screening data are available at https://github.com/jianfcpku/escaping-mutants-screening. Processed mutation escape scores reported previously could be downloaded from https://github.com/jianfcpku/SARS-CoV-2-RBD-DMS-broad. We used vdj_GRCh38_alts_ensembl-5.0.0 as the reference of V(D)J alignment, which can be obtained from https://support.10xgenomics.com/single-cell-vdj/software/downloads/latest. IMGT/DomainGapAlign is based on the built-in lastest IMGT antibody database, and we let the “Species” parameter as “Homo sapiens” while kept the others as default.

Cryo-EM density maps have been deposited in the Electron Microscopy Data Bank with accession codes EMD-32728, EMD-32737, EMD-33552. Structural coordinates have been deposited in the Protein Data Bank with accession codes 7WRJ, 7WRY, 7Y0W, 7Y0V.

## Code Availability

Custom scripts for analyses of sequencing data from rVSV-based mutants screening assays are available at https://github.com/jianfcpku/escaping-mutants-screening.

## Ethical Statement

This study was approved by the Ethics Committee of Beijing Ditan Hospital affiliated to Capital Medical University (Ethics committee archiving No. LL-2021-024-02). Informed consent was obtained from all human research participants.

## Acknowledgments

We thank Sino Biological for the technical assistance on mAbs and RBD expression. We thank J. Luo and H. Lv for the help with flow cytometry. We thank Singlomics Biopharmaceuticals for providing technical help on antibody screening. This project is financially supported by the Ministry of Science and Technology of China and Changping Laboratory under program CPL-1233.

## Author contributions

Y.C. and X.S.X designed the study. Y.C., F.J. and X.S.X wrote the manuscript with inputs from all authors. Q.G. proofed the manuscript. Y.C. and F.S. coordinated the expression and characterization of the neutralizing antibodies. J.W., F.J. and Y.C. performed and analyzed the yeast display screening experiments. Y.Y., T.X., P.W., R.A., Yao W. and X.N. performed the neutralizing antibody expression and characterization, including pseudovirus neutralization assays and ELISA. Y.Y., Youchun W. prepared the VSV-based SARS-CoV-2 pseudovirus. F.Y., Yuhang Z. and A.Z. conceived and performed the rVSV-eGFP-SARS-CoV-2-based mutant screening assays. A.Y., Yao W., S.Y., R.A., W.S. performed and analyzed the antigen-specific single B cell VDJ sequencing. S.D., Ying Z., P.L., Z.Z., J.X. performed the structural analyses. Y.H. and W.Y. performed and coordinated the authentic virus neutralization CPE assay. L.B. and C.Q. performed animal experiments. X.H. and R.J. recruited the SARS convalescents.

## Methods

### Antigen-specific cell sorting, V(D)J sequencing, and data analysis

CD27^+^ IgM^-^ B cells cross-binding to SARS-CoV-1 and SARS-CoV-2 RBD were sorted from SARS-CoV-2 vaccinated SARS convalescents with MoFlo Astrios EQ Cell Sorter (Beckman Coulter). Briefly, CD19+ B cells were enriched from PBMC with EasySep™ Human CD19 Positive Selection Kit II (STEMCELL, 17854). CD19^+^ B cells were then stained with FITC anti-human CD19 antibody (BioLegend, 392508), FITC anti-human CD20 antibody (BioLegend, 302304), Brilliant Violet 421™ anti-human CD27 antibody (BioLegend, 302824), PE/Cyanine7 anti-human IgM antibody (BioLegend, 314532), biotinylated Ovalbumin (SinoBiological) conjugated with Brilliant Violet 605™ Streptavidin (BioLegend, 405229), SARS-CoV-1 biotinylated RBD protein (His & AVI Tag) (SinoBiological, 40634-V27H-B) conjugated to PE-streptavidin (BioLegend, 405204), SARS-CoV-2 biotinylated RBD protein (His & AVI Tag) (SinoBiological, 40592-V27H-B) conjugated to APC-streptavidin (BioLegend, 405207). After washed twice, 7-AAD (Invitrogen, 00-6993-50) were added. 7-AAD^-^, CD19/CD20^+^, CD27^+^, IgM^-^, OVA^-^, SARS-CoV-1 RBD^+^, and SARS-CoV-2 RBD^+^ were sorted.

Isolated SARS-CoV-1 and SARS-CoV-2 RBD cross-binding B cells were then subjected to Chromium Next GEM Single Cell V(D)J Reagent Kits v1.1 following the manufacturer’s user guide (10x Genomics, CG000208). Briefly, cells were resuspended to an appropriated concentration after centrifugation. Cells were processed with 10X Chromium Controller to obtain gel beads-in-emulsion (GEMs) and then subjected to reverse transcription (RT). RT products were subjected to clean up, preamplification and purification with SPRIselect Reagent Kit (Beckman Coulter, B23318). Paired V(D)J sequence were enriched with 10X primers. After library preparation, libraries were sequenced by Novaseq 6000 platform running Novaseq 6000 S4 Reagent Kit v1.5 300 cycles (Illumina, 20028312) or NovaSeq XP 4-Lane Kit v1.5 (Illumina, 20043131).

Sequenced V(D)J raw data were mapped to CRCh38 V(D)J reference by Cell Ranger (v6.1.1). We filtered out non-productive contigs and kept the cell with only one heavy chain and one light chain. With IMGT reference, antibody region and gene annotation were performed by IgBlast (v1.17.1). Mutations were identified by the different nucleotide numbers between antibody and corresponding germline reference.

### High-throughput yeast display-based mutation escape profile

Mutation escape profile assay was performed based on the previously constructed deep mutational scanning libraries of SARS-CoV-2 RBD^6,23^.

Briefly, yeast display libraries were first thawed and amplified in SD-CAA media overnight. Then, RBD expression was induced in SG-CAA media at room temperature with mild agitation for 16-18h. The induced libraries were washed with PBST buffer (PBS with 0.02% Tween-20) and ready for further magnetic-activated cell sorting (MACS)-based mutation escape profiling. Protein A magnetic beads (Thermo Fisher, 10008D) were first conjugated with antibodies by 30min incubation at room temperature, then the Protein A-antibody conjugated beads were washed and incubated with prepared yeast libraries. To obtain pure result, two round of sequential above selections were conducted. The third-round selection was performed using the anti-c-Myc magnetic beads (Thermo Fisher, 88843) to bring down background noise. Yeast cells finally obtained were grown overnight with SD-CAA media and proceed to plasmid extraction with 96 Well Plate Yeast Plasmid Preps Kit (Coolaber, PE053). The extracted plasmid products were used as the template for PCR amplification to detect N26 barcodes as described in^43^. Final PCR products were further purified and submitted to Illumina Next 550 sequencing.

All sequencing data of yeast display deep mutational scanning are processed using the previously described pipeline^23^. In brief, PacBio SMRT reads were mapped to a reference sequence containing SARS-CoV-2 RBD sequence, constant spacer region, and barcode N26, using the Python package dms_variants (v0.8.9), to construct the barcode-variant map. Sequenced barcodes from antibody-selected samples and their corresponding reference samples were also parsed using dms_variants (v0.8.9). Escape scores of each barcode X were defined as F×(n _X,ab_/N_ab_)/(n_X,ref_/N_ref_), where n and N are number of detected barcode X and total barcodes in antibody-selected (ab) or reference (ref) samples, respectively, and F is a scale factor to normalize the scores to 0-1 range. Escape scores of each substitution on RBD were estimated using epistasis models as described previously. For each antibody, only the mutations or residues whose escape scores were at least two times higher than the median score of all mutations or residues were retained. Logo plots for visualization of escape maps were generated using the R package ggseqlogo (v0.1).

### Protein expression and purification for structural analysis

The SARS-CoV-2 S6P (F817P, A892P, A899P, A942P, K986P, and V987P) expression construct that encodes the spike ectodomain (residues 1-1208) with a “GSAS” substitution at the furin cleavage site (residues 682–685) was previously described^21^. Delta mutations (T19R, G142D, 156del, 157del, R158G, L452R, T478K, D614G, P681R, D950N) and Omicron BA.1 mutations (A67V, H69del, V70del, T95I, G142D, V143del, Y144del, Y145del, N211del, L212I, ins214EPE, G339D, S371L, S373P, S375F, K417N, N440K, G446S, S477N, T478K, E484A, Q493R, G496S, Q498R, N501Y, Y505H, T547K, D614G, H655Y, N679K, P681H, N764K, D796Y, N856K, Q954H, N969K, L981F) were further engineered based on this construct. These plasmids were transiently transfected into HEK293F cells using polyethylenimine (Polysciences) to express the corresponding proteins. Culture supernatants were harvested at 96 h post-transfection, concentrated, and exchanged into the binding buffer (25 mM Tris-HCl, pH 8.0, 200 mM NaCl). The proteins were first isolated using the Ni-NTA affinity method, and then further purified using a Superose 6 increase column in the final buffer (20 mM HEPES, pH 7.2, 150 mM NaCl). The antibody Fabs were expressed and purified as previously described^21^.

### Cryo-EM data collection, processing, and structure building

Four microliter S6P protein (0.9 mg/mL) was mixed with the same volume of indicated Fabs (1 mg/mL each), and the resulting mixtures were applied onto glow-discharged Quantifoil holey carbon Au R1.2/1.3 grids using a Vitrobot (Mark IV), at 4 °C and 100% humidity^36^. After ∼3 s, the grids were plunged into the liquid ethane for vitrification. Data were collected using a Titan Krios (operating at 300 kV) equipped with a K3 direct detection camera (Gatan), and processed using cryoSPARC^44^. To improve the local density for the Fab/RBD interface, UCSF Chimera^45^ and Relion^46^ were used to generate masks, and local refinements were further performed using cryoSPARC. COOT^47^ and PHENIX^48^ were used for structural modeling and the real-space refinement, respectively. Structural figures were generated using USCF ChimeraX^49^ and Pymol (Schrödinger, LLC.).

### Crystallization and structure determination

The BD55-1403/BA.1 RBD and BD55-5549/BA.1 RBD complexes were formed by mixing the corresponding protein components at equimolar ratios and incubated on ice for 2 h. The resulting complexes were then further purified using a Superose 200 increase column in the final buffer. Purified Fab/RBD complexes were concentrated to ∼8 mg/mL and subjected to crystal screens. The BD55-1403/BA.1 RBD crystals were obtained by the vapor diffusion method at 18°C, and the reservoir solution contains 2% v/v Tacsimate (pH 5.0), 0.1 M Sodium citrate tribasic dihydrate (pH 5.6), and 16% w/v Polyethylene glycol 3,350. The BD55-5549/BA.1 RBD crystals were obtained in 0.1 M HEPES (pH 7.25), 10% w/v 2-Propanol, and 18% w/v Polyethylene glycol 4,000. For cryo-protection, crystals were soaked in the reservoir solutions supplemented with 17-19% v/v ethylene glycol, and then flash-cooled in liquid nitrogen. Diffraction data were collected at the Shanghai Synchrotron Radiation Facility (beamline BL10U2), and processed then using HKL2000 (HKL Research). The structures were solved by molecular replacement using PHASER^50^, adjusted in COOT and refined using PHENIX.

### Pseudotyped virus neutralization assay

SARS-CoV-2 Spike pseudotyped virus was prepared based on a vesicular stomatitis virus (VSV) pseudotyped virus packaging system as previously reported^6^. Pseudovirus neutralization assays were performed using the Huh-7 cell line (Japanese Collection of Research Bioresources [JCRB], 0403) (for D614G, BA.1, BA.1.1, BA.2, BA.3, BA.2.12.1, BA.4/BA.5, and SARS-CoV-1 pseudotyped virus neutralization assay) or 293T cells overexpressing human angiotensin-converting enzyme 2 (293T-hACE2) (Sino Biological Company) (for Pangolin-GD and RaTG13 pseudotyped virus neutralization assay). Antibodies were serially diluted in DMEM (Hyclone, SH30243.01) and mixed with pseudotyped virus and incubated for 1 h in a 37°C incubator with 5% CO _2_. Digested Huh-7 cells or 293T-hACE2 cells were dispensed to the antibody-virus mixture. After 22-26 hour cells culture in 5% CO_2_, 37 °C incubator, the supernatant was discarded and D-luciferin reagent (PerkinElmer, 6066769) was added, and plates were incubated in darkness for 2 min to allow complete cell lysis. Lysis was transferred to chemiluminescence detection plate and the luminescence value was detected with a microplate spectrophotometer (PerkinElmer, Ensight, 6005290). IC50 was determined by a four-parameter logistic regression model.

### Authentic virus neutralization CPE assay

Neutralization assay for authentic SARS-CoV-2 and its mutant strains using a cytopathic effect (CPE) assay. Monoclonal antibodies or plasma samples were serially diluted in DMEM, mixed with the same volume of the virus, and incubated in a 37°C incubator with 5% CO _2_. The mixture was added to a monolayer of Vero cells (ATCC, CCL-81) in a 96-well plate and cultured for 5 days. All wells were examined under a microscope and the CPE effect of each well was recorded. All experiments were performed in a biosafety level 3 (BSL-3) facility.

### Generation of replication-competent rVSV-SARS-CoV-2-S^BA.1^

Replication-competent rVSV-SARS-CoV-2-S^BA.1^ was generated as described previously, expressing the eGFP reporter gene and the SARS-CoV-2 BA.1 strain spike protein^39,40^. Briefly, 293T cells were transfected with the rVSV backbone plasmid and five support plasmids encoding the T7 polymerase, N, P, M, and L of VSV using the calcium phosphate method. Recovery of the virus was determined by cytopathic effects and eGFP expression. Viruses in the supernatant were harvested and passaged on Vero cells to obtain viral stocks.

### Selection of escape mutations

The rVSV-SARS-CoV-2-S^BA.1^ mutants that evade mAb neutralization were screened according to a previous report. Briefly, rVSV-SARS-CoV-2-S^BA.1^ (5 × 10^5^ FFU) was preincubated with 5-fold serial dilution of monoclonal antibody at RT for 30 min. Then the antibody-virus mixtures were added to monolayers of Vero cells and incubated at 37°C. After four days, the supernatants from the wells containing the highest concentration of mAbs that showed extensive GFP-positive foci were harvested. For the next round of selection, the supernatants were passaged on fresh Vero cells in the presence of the same concentrations of mAbs as before. The passages continued until 50 μg/ml mAbs could no longer neutralize the mutant viruses.

Viral RNA was extracted from the supernatants of each passage using the QIAamp Viral RNA Mini Kit (Qiagen, US). Purified RNAs were reverse transcribed using PrimeScript RT reagent Kit with gDNA Eraser (Takara, JP) and the S gene was amplified using PrimeSTAR GXL DNA Polymerase (Takara, JP) for next-generation sequencing.

Viral RNA was extracted from the supernatants of each passage using the QIAamp Viral RNA Mini Kit (Qiagen, US). Purified RNAs were reverse transcribed using PrimeScript RT reagent Kit with gDNA Eraser (Takara, JP) and the S gene was amplified using PrimeSTAR GXL DNA Polymerase (Takara, JP) for next-generation sequencing.

### Identification of enriched escape substitutions

Illumina sequencing adapters were first trimmed from raw sequencing reads using trim_galore (default parameters, version 0.6.7, cutadapt version 1.18). Trimmed reads were mapped to the nucleotide sequence of SARS-CoV-2 BA.1 spike glycoprotein (codon-optimized for rVSV) using hisat2 (default parameters, v.2.2.1). PCR duplicates were removed from mapped reads using Picard MarkDuplicates (REMOVE_DUPLICATES=TRUE, version 2.18.29). SNP information was extracted from deduplicated BAM files using bcftools mpileup (-d 900000 - C 50 -Q 30 -q 5, version 1.8). Detected nucleotide mutations at each position were then parsed using custom Python scripts to remove all insertions or deletions, and converted into the occurrence ratio of residue substitutions.

### In vivo SARS-CoV-2^BA.1^ challenge in mice

The mice used in this study were 6-8 weeks old. Murine studies were performed in an animal biosafety level 3 (ABSL3) facility using HEPA-filtered isolators. All procedures in this study involving animals were reviewed and approved by Institutional Animal Care and Use Committee of the Institute of Laboratory Animal Science, Peking Union Medical College (BLL22001). Mice were inoculated intranasally with authentic SARS-CoV-2 BA.1 (SARS-CoV-2/human/CHN/Omicron-1/2021 (Genbank: OM095411.1)) stock virus at 1 × 10^5^ TCID50. The infected mice were observed daily to record body weights and were sacrificed at 3 days post infection, and the lungs and tracheas were collected for viral load detection.

Viral load analysis was performed by qRT-PCR. The total RNA of the lungs and trachea was extracted with the RNeasy Mini Kit (Qiagen). Lung and trachea homogenates were prepared by using an electric homogenizer. The reverse transcription was processed with PrimerScript RT Reagent Kit (TaKaRa) according to the manufacturers’ instructions. qRT-PCR reactions were performed using the PowerUp SYBG Green Master Mix Kit (Applied Biosystems), according to following cycling protocol: 50 °C for 2 min, 95 °C for 2 min, followed by 40 cycles at 95 °C for 15 s and 60 °C for 30 s, and then 95 °C for 15 s, 60 °C for 1 min, 95 °C for 45 s. Forward primer 5′-TCGTTTCGGAAGAGACAGGT-3′ and reverse primer 5′-GCGCAGTAAGGATGGCTAGT-3 ′ were used in qRT-PCR. Standard curves were constructed by using 10-fold serial dilutions of recombinant plasmids with known copy numbers (from 1.47 × 10^9^ to 1.47 × 10^1^ copies/μl).

## Figure Legends

**Fig. 1** | **Identification of broad sarbecovirus neutralizing antibodies**.

**a**, Schematic for the identification and characterization of bsNAbs, and selection of bsNAb drug candidates from SARS-CoV-2-vaccinated SARS convalescents. **b**, Neutralizing IC50 against SARS-CoV-2 variants VSV-based pseudoviruses and authentic viruses, and ELISA EC50 against sarbecovirus RBD by identified bsNAbs and other published antibodies in group E1, E3 and F3.

**Fig. 2** | **Proposed strategies for non-overlapping bsNAb combination**

**a-b**, Aligned structures of BD55-3152 (Group E1, PDB: 7WR8) together with **a**, BD55-1239 (Group F2, PDB: 7WRL) or **b**, BD55-3372 (Group F3, PDB: 7WRO) in complex with SARS-CoV-2 BA.1 RBD. **c**, Aligned structure shows BD55-1239 (PDB: 7WRL) in F2 and BD55-3372 (PDB: 7WRO) in F3 could not form non-overlapping cocktails.

**Fig. 3** | **Escape maps and structural analyses of representative E1 and F3 antibodies**.

**a-b**, Average escape scores of antibodies in group E1 (a) and F3 (b), and escape maps of selected drug-candidate bsNAbs in group E1 (a) and F3 (b). MSA on the corresponding escaping hotspot residues are shown. **c**, Structure of BD55-3546 bound to RBD_Delta_. **d**, Structure of BD55-5549 bound to RBD_BA.1_. **e**, Structure of BD55-4637 bound to RBD_BA.1_. **f**, Structure of BD55-5514 and BD55-5840 bound together to RBD_BA.1_. **g**, Interactions between BD55-5514 and RBD_BA.1_. **h**, Interactions between BD55-5840 and RBD_BA.1_.

**Fig. 4** | **SA55/SA58 cocktail robustly neutralizes SARS-CoV-2 escaping mutants**.

**a**, Schematic of in vitro escape mutation screening using SARS-CoV-2^BA.1^-spike-pseudotyped replication-competent recombinant VSVs (rVSV). The virus was passed in the presence of antibody dilutions for 4 days on Vero cells. Cells were screened for virus replication by monitoring GFP^+^. **b**, Antibody concentration of dilutions that are escaped, passed, and sequenced of each passage and each antibody. SA55 and SA58 alone were tested in two biological replicates, respectively. SA55+SA58 cocktail was tested in four biological replicates. **c-e**, The ratio of enriched escape mutations of each replicate and each passage by deep sequencing. **c**, using SA58 only; **d**, using SA55 only; **e**, SA55+SA58 cocktail. **f**, Neutralizing activities by SA55, SA58 and SA55+SA58, against selected SARS-CoV-2 variants harbouring potential escaping mutations of SA55 or SA58. **g**, Ratio of selected SA55/SA58-escaping mutations in detected SARS-CoV-2 sequences from GISAID.

**Fig. 5** | **SA55/SA58 demonstrate potent protection efficacy in mice**.

**a-b**, Design for the SARS-CoV-2 BA.1 challenge experiment in mice. In prophylactic groups, 20mg/kg SA55+SA58 were administered intraperitoneally or intramuscularly and challenged by SARS-CoV-2 BA.1 intranasally one day later. In therapeutic groups, antibodies were given intraperitoneally 2 hours after virus challenge. Each group consists of 5 mice. **c**, Percentage of changes in body weight compared to the weight of each mouse before the experiment. Error bars indicate mean±s.d. **d**, Viral load in lung (up) and trachea (bottom) of mice in each group. Each point corresponds to samples from a mouse. P-values were calculated using one-tailed Wilcoxon rank-sum test. The alternative hypothesis is that the values in prophylactic or therapeutic groups are less than values in control groups.

## Extended Data Figures

**Extended Data Fig. 1** | **FACS strategy to isolate cross-reactive memory B cells from SARS-CoV-2-vaccinated SARS convalescent plasma**.

**Extended Data Fig. 2** | **Structural models of representative broadly reactive antibodies**. Aligned structures of representative antibodies in epitope groups with broad reactivity against sarbecoviruses are shown, including S309 in E1 (PDB: 6WPS), S2H97 in E3 (PDB: 7M7W), S304 in F1 (PDB: 7JW0), BD55-1239 in F2 (PDB: 7WRL), and ADG-2 in F3 (PDB: 7U2D).

**Extended Data Fig. 3** | **Detailed information about the 12 bsNAb drug candidates**.

SHM counts and rates in nucleotides and amino acids, in addition to the germline V-J gene combinition of the 12 candidates from 3 epitope groups are shown in the table.

**Extended Data Fig. 4** | **Workflow for the cryo-EM 3D reconstruction**.

**a-c**, Cryo-EM data collection and processing workflow for the reconstruction of the structures of **a**, BD55-3546/Delta S6P complex; **b**, BD55-4637/BA.1 S6P complex; **c**, BD55-5514+BD55-5840/BA.1 S6P complex.

**Extended Data Fig. 5** | **Detailed interactions between Group E1 and F3 bsNAbs and SARS-CoV-2 RBD**.

**a**, Interactions between BD55-3546 and SARS-CoV-2 Delta RBD. **b**, Interactions between BD55-5549 and SARS-CoV-2 BA.1 RBD. **c**, Interactions between BD55-4637 and SARS-CoV-2 BA.1 RBD.

**Extended Data Fig. 6** | **Identification of non-overlapping antibody cocktails**.

Cryo-EM structure of SA55+SA58 Fab in complex with BA.1 S6P. SA55 binds “up” RBD only, while SA58 binds both “up” and “down” RBD.

**Extended Data Fig. 7** | **Psuedovirus neutralizing IC50 of SA55 and SA58 against Omicron variants harboring selected single substitutions**.

IC50 fold changes against constructed pseudoviruses compared to those against BA.2 were shown.

**Supplementary Table 1**. Summarized information of SARS-CoV-2-vaccinated SARS-CoV-1 convalescents.

**Supplementary Table 2**. Neutralizing activities and binding capabilities against sarbecoviruses of 314 broad sarbecovirus neutralizing antibodies.

**Supplementary Table 3**. Crystal data collection, refinement and validation statistics.

**Supplementary Table 4**. Cryo-EM data collection, refinement and validation statistics.

**Extended Data Fig. 1.**
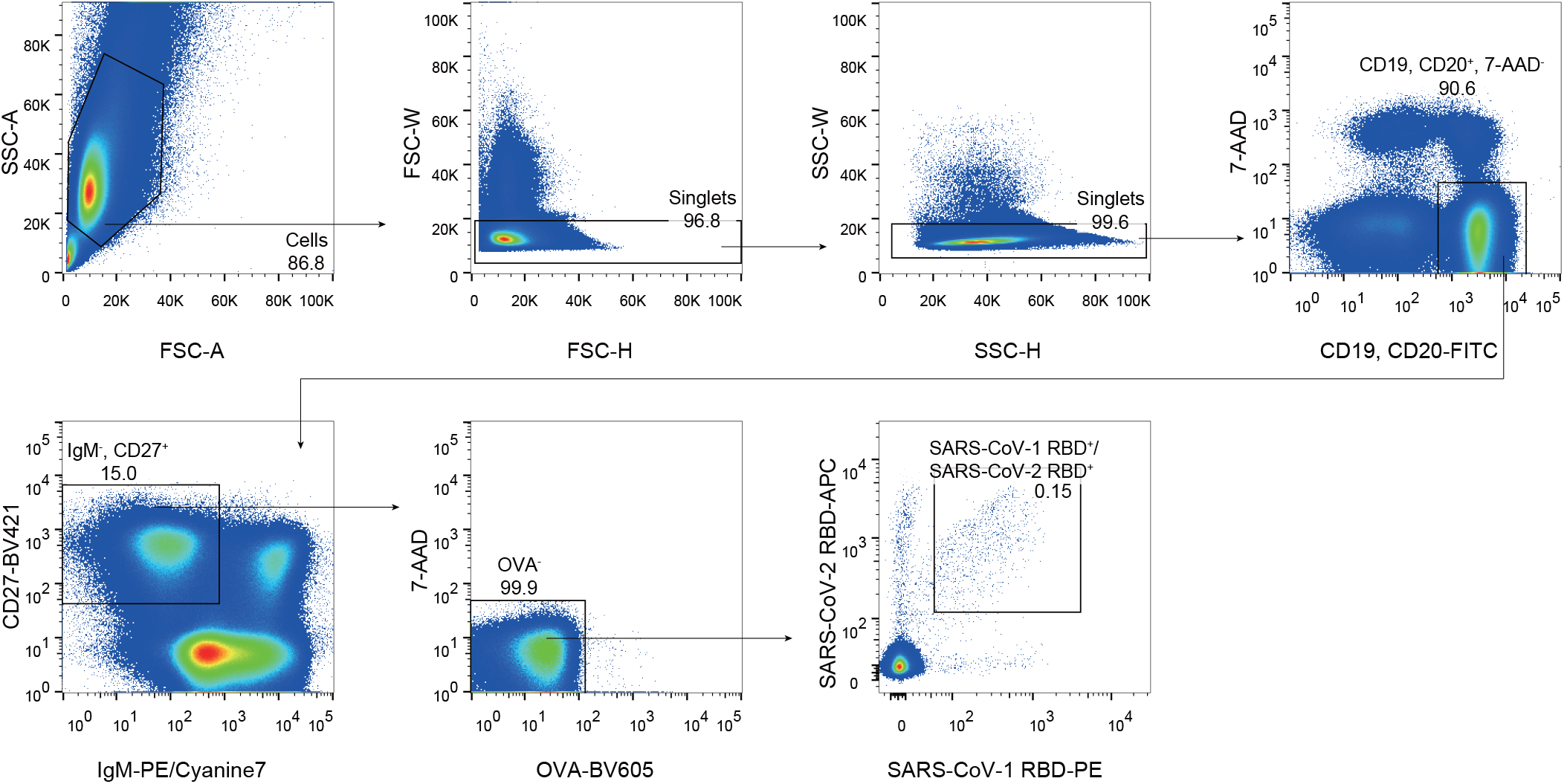
FACS strategy to isolate cross-reactive memory B cells from SARS-CoV-2-vaccinated SARS convalescent plasma.

**Extended Data Fig. 2.**
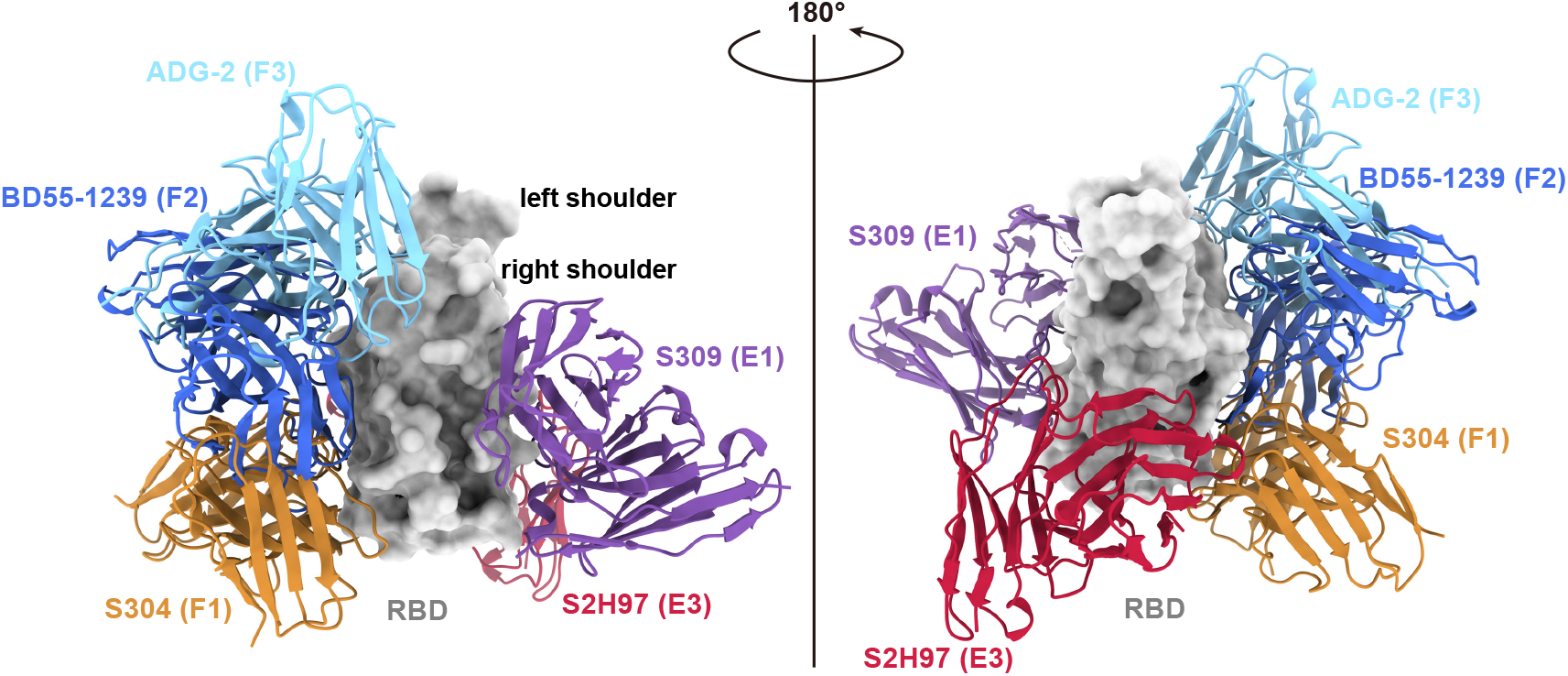
Structural models of representative broadly reactive antibodies. Aligned structures of representative antibodies in epitope groups with broad reactivity against sarbecoviruses are shown, including S309 in E1 (PDB: 6WPS), S2H97 in E3 (PDB: 7M7W), S304 in F1 (PDB: 7JW0), BD55-1239 in F2 (PDB: 7WRL), and ADG-2 in F3 (PDB: 7U2D).

**Extended Data Fig. 3.**
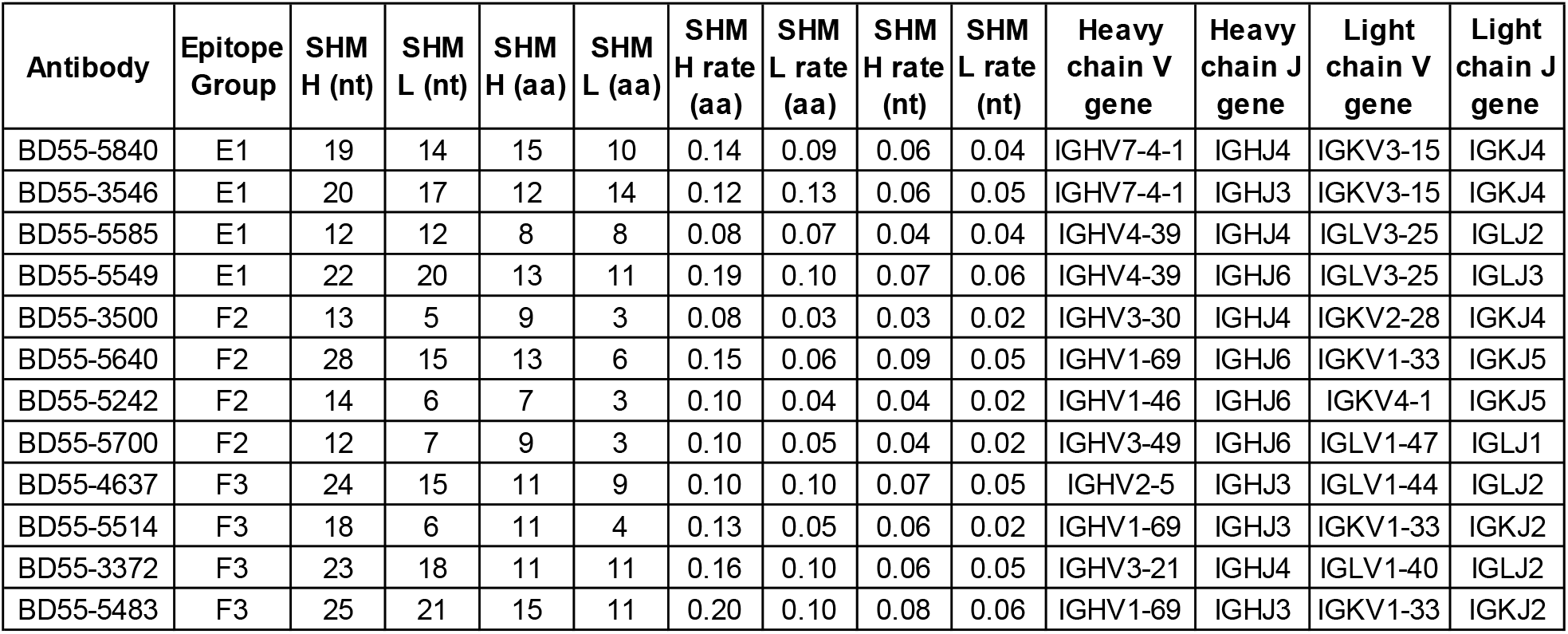
Detailed information about the 12 bsNAb drug candidates. SHM counts and rates in nucleotides and amino acids, in addition to the germline V-J gene combinition of the 12 candidates from 3 epitope groups are shown in the table.

**Extended Data Fig. 4.**
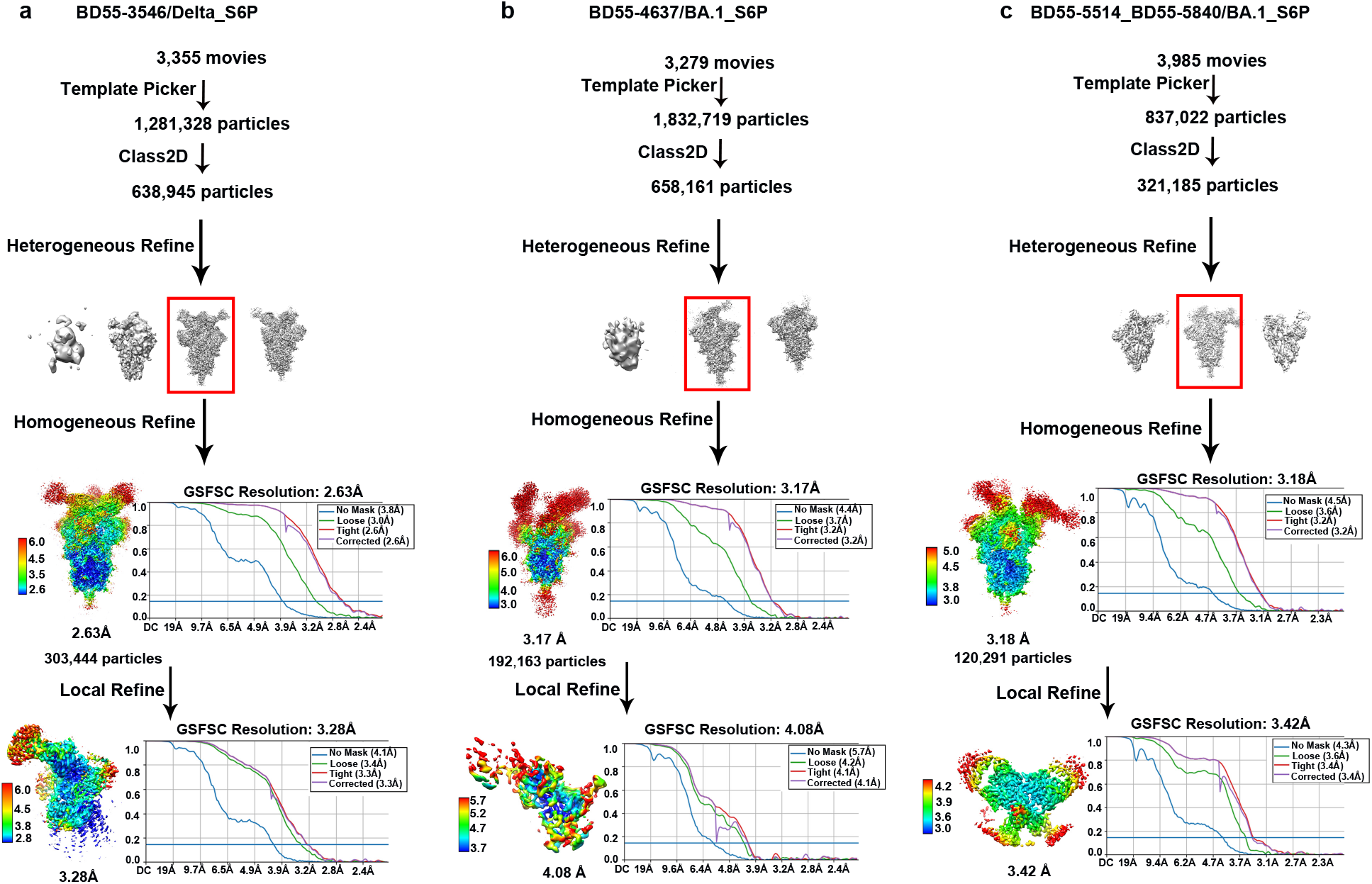
Workflow for the cryo-EM 3D reconstruction. a-c, Cryo-EM data collection and processing workflow for the reconstruction of the structures of a, BD55-3546/Delta S6P complex; b, BD55-4637/BA.1 S6P complex; c, BD55-5514+ BD55-5840 (SA55+SA58)/BA.1 S6P complex.

**Extended Data Fig. 5.**
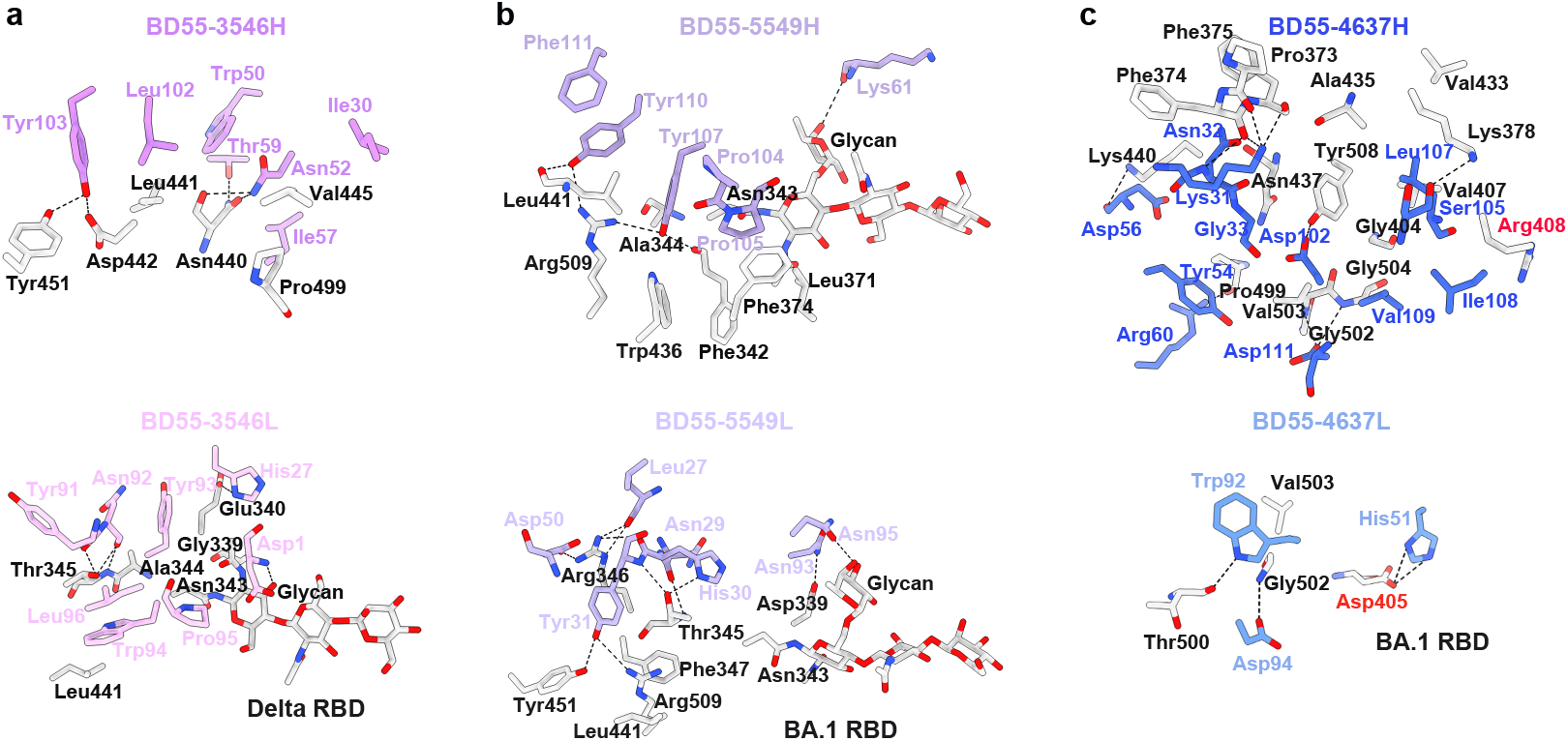
Detailed interactions between Group E1 and F3 bsNAbs and SARS-CoV-2 RBD. a, Interactions between BD55-3546 and SARS-CoV-2 Delta RBD. b, Interactions between BD55-5549 and SARS-CoV-2 BA.1 RBD. c, Interactions between BD55-4637 and SARS-CoV-2 BA.1 RBD.

**Extended Data Fig. 6.**
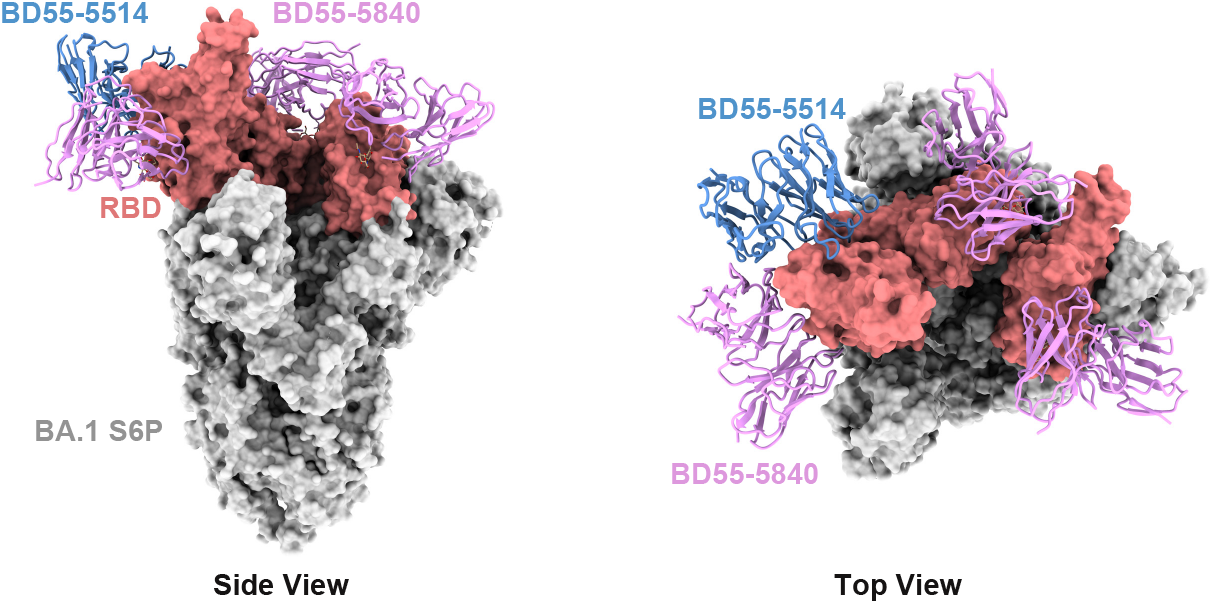
Identification of non-overlapping antibody cocktails. Cryo-EM structure of SA55+SA58 Fab in complex of BA.1 S6P. SA55 binds “up” RBD only, while SA58 bind both “up” and “down” RBD.

**Extended Data Fig. 7.**
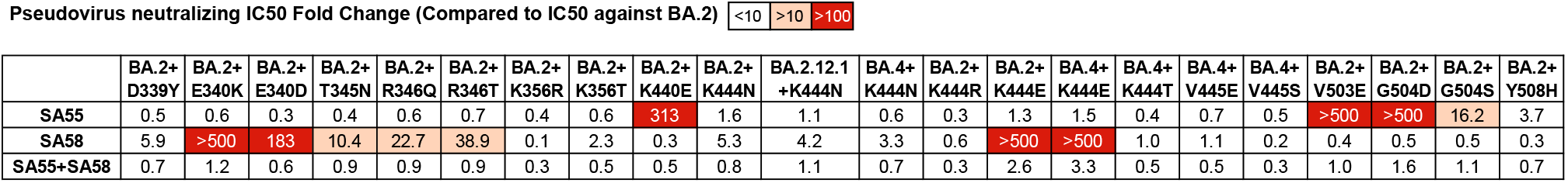
Psuedovirus neutralizing IC50 of SA55 and SA58 against Omicron variants harboring selected single substitutions. IC50 fold changes against constructed pseudoviruses compared to that against BA.2 were shown.

